# An optogenetic assay for the dauer decision in Caenorhabditis elegans

**DOI:** 10.1101/2024.04.15.589657

**Authors:** Ariel Parker, James Mullins, Abel Corver, Anastasia Miller, Iman Mosley, Andrew Gordus

**Affiliations:** Department of Biology, Johns Hopkins University, Baltimore, MD; Solomon H. Snyder Department of Neuroscience, Johns Hopkins University, Baltimore, MD; Department of Biology, Lund University, Lund, Sweden

## Abstract

The dauer decision in *Caenorhabditis elegans* is a critical developmental decision that ensures survival under harsh environmental conditions. Factors such as temperature, food availability, and pheromone strongly influence the decision to enter and exit dauer. Traditional assays that assess the dauer decision are often confounded by the influence of pheromones from the population, which are often dynamic and highly variable. To mitigate this issue, we developed a simple, single-housing assay for dauer quantification that is compatible with optogenetics. We show that insulin-like peptides (ILPs) from ASJ and other neurons strongly influence the decision to exit dauer, and that ASJ activity can be manipulated with optogenetics to influence the dauer decision in a temporally precise manner.

## Introduction

Dauer is an optional environmental stress-response stage that replaces the third larval stage (Figure 1A). Compared to conspecifics of similar age, dauer worms are thinner, do not eat, thicken their outer cuticle, seal their mouth and anus, and show several other anatomical differences. These changes increase the worm’s chances of survival under harsh environmental conditions: a thickened cuticle prevents desiccation while sealing orifices prevents eating and defecation, both of which are metabolically active activities.

**Figure 1:**
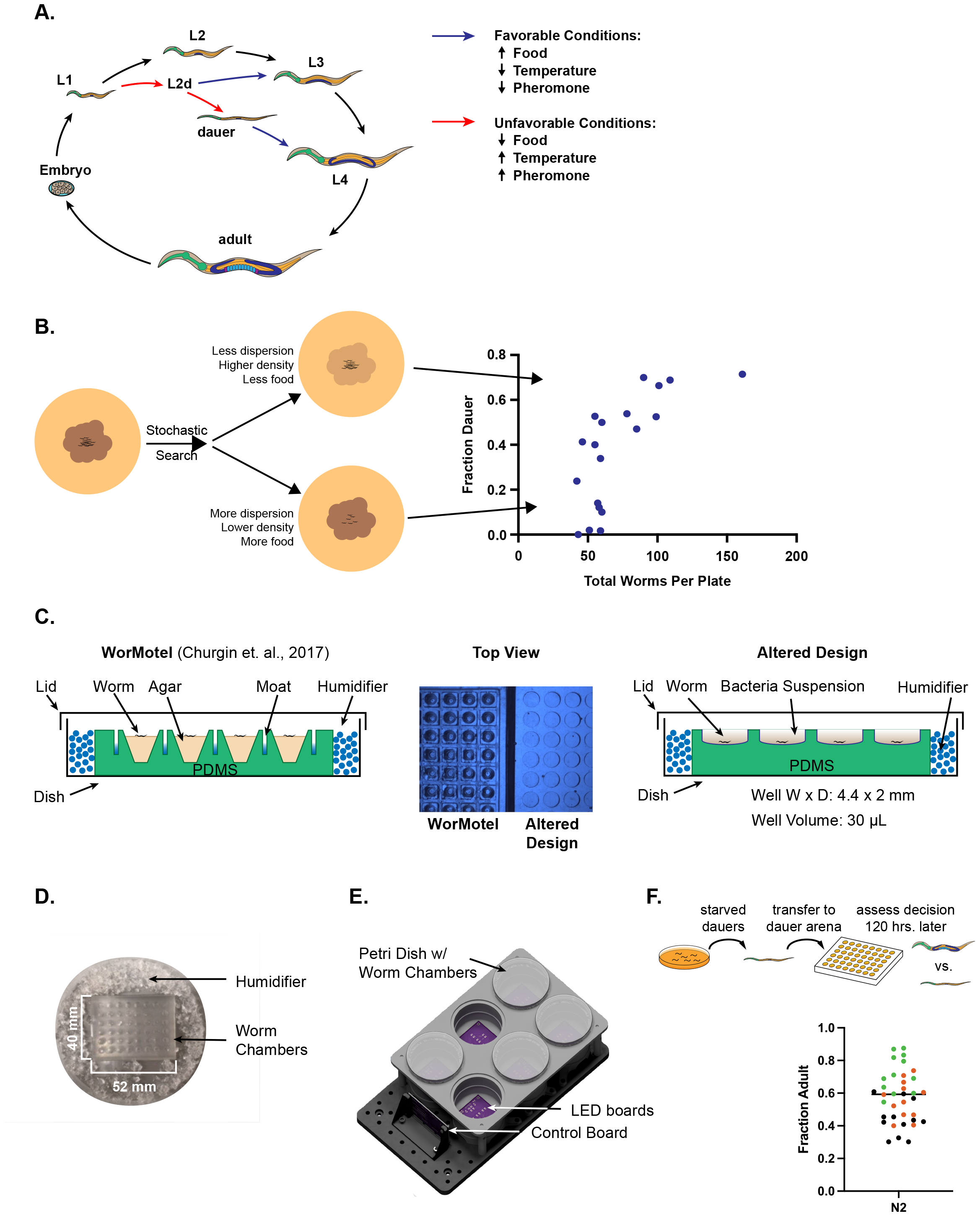
Dauer Assays and the Dauer-WorMotel. **A**. *C. elegans* life cycle. Worms proceed through four larval stages until reproductive adulthood. After L1, worms choose to enter L2d (a pre-dauer stage) and subsequently choose to enter dauer. Once under dauer arrest, worms monitor the external environment for improving conditions as a sign to proceed with development into L4. **B**. Plate-based dauer assay. Fraction dauer worms versus the number of worms on the plate, from dauer entry experiments on petri dishes. Five adult worms laid eggs on dauer assay plates for 4 hours were removed, and the plates were stored at 25 °C for 8 days (Neal et al., 2013). After 8 days, the number of dauers and non-dauer worms were counted. Each dot represents the fraction of dauer worms on a single petri dish. Data were collected on the same day. **C**. The WorMotel and the Dauer-WorMotel. The WorMotel (left) was designed to individually house worms for longevity assays. The Dauer-WorMotel (right) relies on liquid bacterial suspension that replaces the agar-food bedding of the original design. The original WorMotel has conical wells, while the Dauer-WorMotel has flat-bottomed wells that improve visibility. **D**. Humidified Chamber: To prevent desiccation of the small (23 uL) volumes, the worm chamber is stored in a 10 cm petri dish surrounded by AgSap® hydroscopic crystals that serve as a humidifier. **E**. Optogenetics set-up for dauer exit assay. A single Teensy 3.2 controls the lights for up to 6 Dauer-WorMotels. **F**. Dauer exit assay. Dauer worms are harvested after starvation and SDS selection, and individually housed in separate wells of the Dauer-WorMotel in 23 µL of 1 mg/mL live HB101 (with 5 mg/mL cholesterol) at 20 °C. N2 worms exit dauer under the experimental conditions of this paradigm. Dauers versus L4/adults were counted after the five days. Each dot represents one Dauer-WorMotel, which contains up to 48 dauer worms (L4/adult average = 0.58, standard deviation = 0.16, n = 1617 individual worms). The different colors represent 3 different experimental days (12 Dauer-WorMotels per day).

Changes to neuronal morphology in the chemosensory and mechanosensory organs also occur (Albert & Riddle, 1983). For example, the nerve endings of the ASI neuron (known to be important for dauer entry (C I. Bargmann & Horvitz, 1991)) are shortened and displaced from the bundle of other amphid organ neuronal endings (Albert & Riddle, 1983). Alternatively, some neuron’s sensation capabilities are strengthened during dauer. AWC, an important chemosensory neuron, has large cilia in the dauer stage versus non-dauers (Albert & Riddle, 1983).

Temperature, food availability, and population density sensed via pheromones are the three most salient environmental variables that influence dauer entry and exit (Ailion & Thomas, 2000; Butcher *et al*., 2007). Neurons in chemosensory organs like the amphid organ sense odorants, pheromones, and food signals. ASI is a polymodal neuron that senses ascaroside pheromones (Park *et al*., 2012) and a shortened range of temperature stimuli (Beverly *et al*., 2011). ASJ senses bacterial food metabolites to promote dauer exit (Kaul et al., 2014a).

Neuroendocrine signaling strongly influences the dauer decision. Various insulin-like peptides work either as agonists (like INS-6 and INS-28) or as antagonists (like INS-7 and INS-15) on DAF-2, the sole insulin-like growth factor receptor (IGFR) in the worm, to respectively reduce or promote dauer formation (Zheng *et al*., 2018). A high ratio of agonists to antagonists is maintained by good environmental conditions, which increases DAF-2 activity in the intestine and neurons (Hung *et al*., 2014). The TGF-β-related ligand DAF-7 is known to promote reduce dauer formation and support continuous development, most likely due to its role as a signal of satiety; dauer-inducing conditions reduce *daf-7* expression in the ASI chemosensory neurons (Ren *et al*., 1996).

Traditional dauer assays use petri dishes to assess animal decisions in bulk (Golden & Riddle, 1984; Neal *et al*., 2013). To provide unfavorable conditions, petri dishes containing synthesized dauer-inducing ascarosides are seeded with limited amounts of bacterial food source. Eggs are laid on these plates and incubated under warm temperatures (25 °C). Subsequently, the number of dauer versus non-dauer worms are counted 72 to 84 hours later. A significant challenge with this approach is that is performed on a population of worms, but the dauer decision is strongly influenced by the pheromone of neighboring animals.

There were two challenges with the petri-dish assay: variable life history and variable population density. When food is plentiful, the worms rarely leave the food patch (Milward *et al*., 2011). However, as food concentration decreases, worms become more exploratory, and are more likely to leave the bacteria lawn. This produces a more variable sensory and metabolic history for each worm on the plate, even though all worms were placed under “the same” conditions. Environmental food variability is encoded neuronally and is known to affect worm behavior and decision-making (Calhoun *et al*., 2015).

Greater exploration also leads to a greater number of worms leaving the agar and dying on the sides of the petri dish, which decreases the amount of ascarosides across the assay plates and increases food availability. Both food and ascarosides influence the dauer decision. Since worm exploration is more variable under dauer-inducing conditions, the number of animals that die on the sides of the plate is also more variable, which in turn produces more variable food and pheromone conditions during the timescale of the experiment. In prior work (Neals et al) this confound has been mitigated by only quantifying data from plates that had 65 – 85 worms.

However, this approach requires ignoring a large fraction of the experiments which did not fall within this narrow range of conditions.

To mitigate this issue, recent advances have been made in controlling the environment of populations of animals in bulk or individually using microfluidics. However, these systems are not trivial to implement, and require constant monitoring to prevent biofilm build-up. Non-microfluidic systems to monitor individual worm development have been developed, but these are not amenable to the low food conditions that are typical for dauer assays.

Here, we develop a simple microwell assay that is amenable to optogenetics. We test dauer exit with this assay under a variety of genotypes that perturb insulin-like growth factor (IGF) signaling, and manipulate the dauer exit decision via optogenetic stimulation of the sensory neuron, ASJ. We find 12 hours of ASJ stimulation is sufficient to induce dauer exit, but 6 hours is insufficient.

## Materials and Methods

### Worm strains

Wild-type and mutant strains used originate from the Bristol strain N2 (S Brenner, 1972). Strains were cultivated, optionally in the presence of all-*trans* retinal (Sigma-Aldrich), as described previously (JF Liewald *et al*, 2008). Strains used were: **N2** (laboratory strain), **ZM6523** *ins-4;ins-5;ins-6* (*hpDf761*) and **CB1370** *daf-2* (*e1370*), **AGG0119** *Ptrx-1::TeTx::mCherry* (*zucEx0109*), **AGG0127** *Ptrx-1::TeTx::mCherry; ins-4;ins-5;ins-6* (zucEx0117), **AGG0136** *Ptrx-1::TeTx::mCherry; daf-2 (e1307)* (zucEx0127), **AGG0122** *Ptrx-1::ChR(C128S)::GFP* (*zucEx0112*). **ZM6523** and **CB1370** were obtained from the Caenorhabditis Genetics Center.

### The dauer exit assay for N2 worms

Agar plates (10 cm) were seeded with 900 μL HB101 *E. coli* and stored at 20 °C for two days. Twelve gravid adult N2 worms were picked on to these plates, and after one week, SDS selection and sucrose flotation were used to isolate dauers (Karp, 2018). Dauer worms were then individually pipetted into wells of the Dauer-WorMotel, pre-filled with 23 μL 1 mg/mL live *E. coli* HB101 in S-Medium supplemented with 5 mg/mL cholesterol. The Dauer-WorMotels were sealed in petri dishes with hydrated AgSap® hydroscopic crystals (which serve as a humidifier for the assay, M^2^ Polymer Technologies) and placed at 16.9 – 17.3 °C. After 5 days (120 hours) dauers, L4s and adults were counted.

### The dauer exit assay for daf-2 worms

Agar plates (10 cm) were seeded with 900 μL HB101 *E. coli* and stored at 20 °C for two days. Twelve gravid adult *daf-2* worms were picked on to these plates and stored at 20 °C for 24 hours. The next day, plates were transferred to 25 °C for three additional days to induce dauer formation. Dauer worms were then individually pipetted into wells of the Dauer-WorMotel, pre-filled with 23 μL live *E. coli* HB101 (OD_600_ = 0.27-0.32) in S-Medium supplemented with 5 mg/mL cholesterol. The Dauer-WorMotels were sealed in petri dishes with hydrated hydroscopic crystals and placed at 16.9 – 17.3 °C. After 4 days (96 hours) dauers, L4s and adults were counted.

### LED and control board setup

We built a custom LED circuit board to control the lights need to excite the cation channelrhodopsin in the ASJ neuron. Blue and amber LED intensities were controlled by pulse-width modulation (PWM) at 1 MHz frequency, which is much faster than neuronal response properties. PWM timing was controlled by a Teensy 3.2. Light intensities at different PWM settings were calibrated using a Compact Power and Energy Meter Console with Digital 4” LCD (PM100D, Thor Labs) and Standard Photodiode Power Sensor (S121C, Thor Labs).

### Optogenetic dauer exit assay

Agar plates (10 cm) were seeded with 900 μL 500 μM trans-retinal in HB101 *E. coli* and stored in the dark at 20 °C for two days. Twelve gravid adult worms were picked on to these plates and stored at 20 °C in the dark for 24 hours. The next day, plates were transferred to 25 °C in the dark for three additional days to induce dauer formation. Dauer worms were then individually pipetted into wells of the Dauer-WorMotel, pre-filled with 23 μL live *E. coli* HB101 (OD_600_ = 0.27-0.32) in S-Medium supplemented with 5 mg/mL cholesterol. The Dauer-WorMotels were sealed in petri dishes with hydrated hydroscopic crystals and placed at 16.9 – 17.3 °C on the optogenetics rigs. LEDs were programmed for individual experiments. After 4 days (96 hours), dauers, L4s and adults were counted.

## Results

### 1. The Dauer-WorMotel: a dauer assay with controlled conditions

We developed a standardized assay to study the dauer exit decision by combining elements of previous dauer entry assays with a different means of housing the worms during the experiment. Traditionally, assays to assess dauer decisions were performed on petri dishes, with many eggs/worms initially placed on the plates to be counted later. For the dauer exit assay for this work, we wanted to isolate the worms, to standardize food conditions, make counting easier and to prevent ascaroside signaling between worms.

We modified the WorMotel for this goal. The original WorMotel was developed by the Fang-Yen laboratory for longevity assays (Churgin *et al*., 2017). In their set-up, worms are individually housed in wells with agar bedding and ad libitum bacterial food (Figure 1C). This assay has several advantages. First, individually housed worms are easier to count. Second, this set-up controls population density – an individually housed worm does not sense pheromone from other worms to influence its own decision. Third, the transparency of PDMS is amenable to optogenetic stimulation.

While the original WorMotel design worked well for longevity assays, it did not work well for dauer assays. For the longevity assays, worms were provided with a high density of food. When they left the patch and encountered the copper sulfate moat (an aversive cue), the worms would return to the food patch. However, under the low-food conditions used for dauer assays, the worms were less risk-averse, and nearly all of them ended up in the copper moats within a few hours (data not shown).

To make the WorMotel more amenable for dauer assays, we optimized a variety of features from the original design (Figure 2C). In the original design, copper moats prevented the escaping of worms from the agar bedding; in our design, we removed the copper moats and housed the worms in liquid media medium - a combination of the bacterial food source and S-Medium (Stiernagle, 2006) (Figure 1C-D). The original WorMotel had conical-shaped wells; we amended the wells to be more cylindrical to improve optics so worms could be reliably phenotyped in the wells (Figure 1C-D). With these changes, we refer to the set-up as the Dauer-WorMotel. A feature of the original WorMotel that we did keep was the use of polydimethylsiloxane (PDMS). PDMS is a clear, inert silicone-based polymer used in microfluidics and other biological applications (Raj M & Chakraborty, 2020). It is transparent and compatible with the illumination necessary for optogenetics. We built an array of custom-designed LED boards and holders to house the Dauer-WorMotel, and programmed a Teensy 3.2 to dynamically control illumination. We used this asay in subsequent experiments to quantify dauer exit.

**Figure 2:**
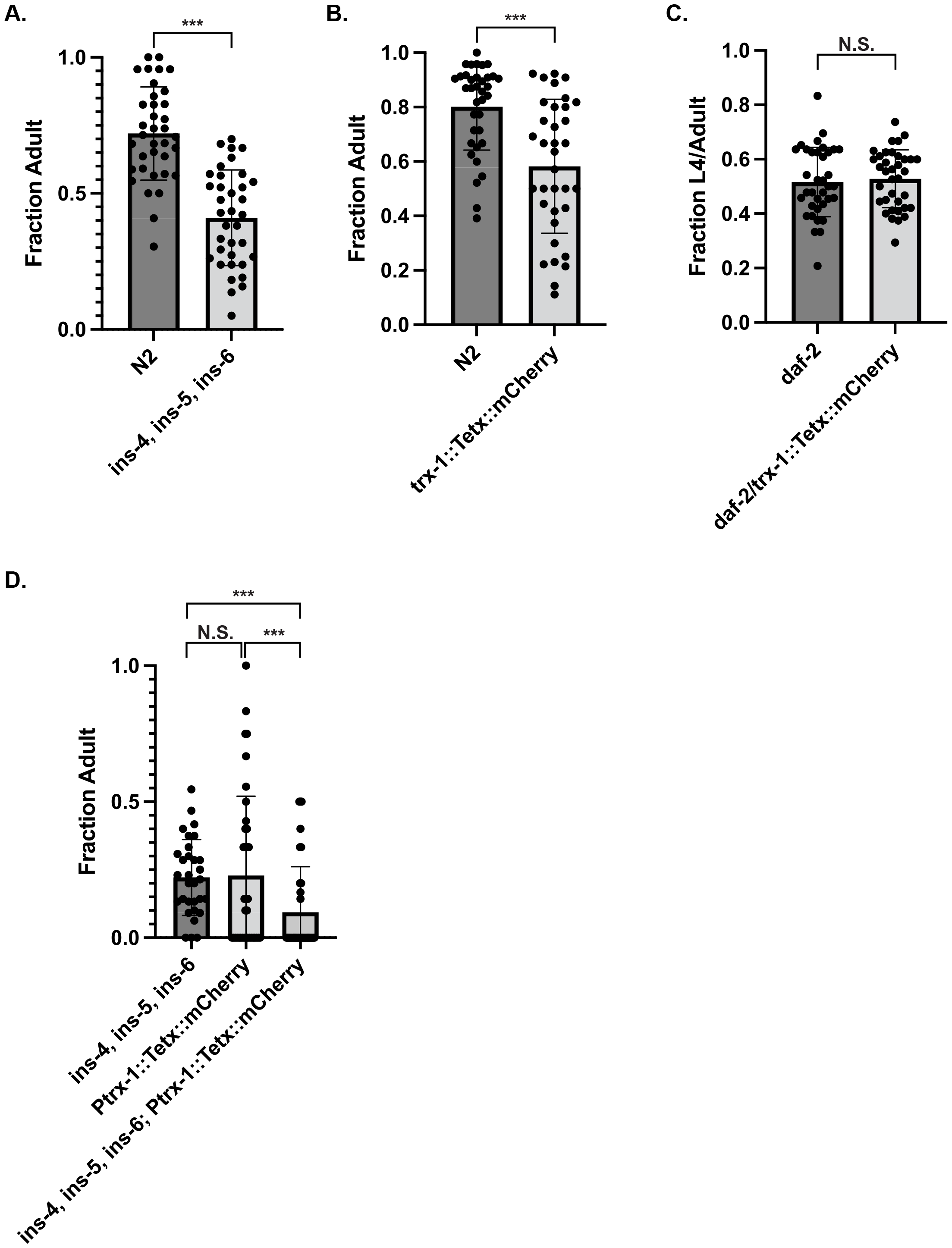
ILP influence of dauer exit in the Dauer-WorMotel. **A**. ASJ ILPs drive dauer exit. Dauer exit was quantified in the N2 and *ins-4;ins-5;ins-6* triple mutant, ZM6523. (X^2^ (1, N = 1567) = 758.0, p <0.001). **B**. Synaptic release from ASJ is influences dauer exit. Dauer exit was quantified in the N2 and *Ptrx-1::TeTx::mCherry* background (X^2^ (1, N = 1134) = 333.0, p <0.001). **C**. The influence of synaptic activity of ASJ on dauer exit relies on DAF-2 activity. Dauer exit was quantified in *daf-2 (e1370)* and *daf-2,Ptrx-1::TeTx::mCherry* (X^2^ (1, N = 1454) = 635.0, p = 0.39). **D**. *ins-4/ins-6* neuromodulation and ASJ chemical neurotransmission work in parallel to influence dauer exit. We observed less dauer exit in the *ins-4/ins-5/ins-6; Ptrx-1::TeTx::mCherry* cross as compared to either single mutant. The chi-square statistics are as follows. Between *ins-4/ins-5/ins-6* and *Ptrx-1::TeTx::mCherry*, X^2^ (1, N = 628) = 232.0, p = 0.35. Between *ins-4/ins-5/ins-6* (ZM6523) and *ins-4/ins-5/ins-6; Ptrx-1::TeTx::mCherry*, X^2^ (1, N = 549) = 153.0, p < 0.001. Between *Ptrx-1::TeTx::mCherry* and *ins-4/ins-5/ins-6; Ptrx-1::TeTx::mCherry*, X^2^ (1, N = 389) = 153.0, p < 0.001.

Dauer animals were produced by starving worms, and using SDS selection to purify dauer animals from the population. Wen then housed these worms in the Dauer-WorMotel under different conditions. In Table 1, we Dauer-WorMotels were loaded with different food amounts and placed at different temperatures to quantify the effects of these variables on dauer exit. Higher temperature consistently inhibited dauer exit, regardless of food concentration (Table 1). Lower temperatures combined with high food availability reliably increased dauer exit. Based on these results, the final conditions for Dauer-WorMotel experiments with N2 worms were – 1 mg/mL live HB101 at 20 °C which produced an average dauer exit probability of 0.58 (Figure 1F).

**Table 1.**
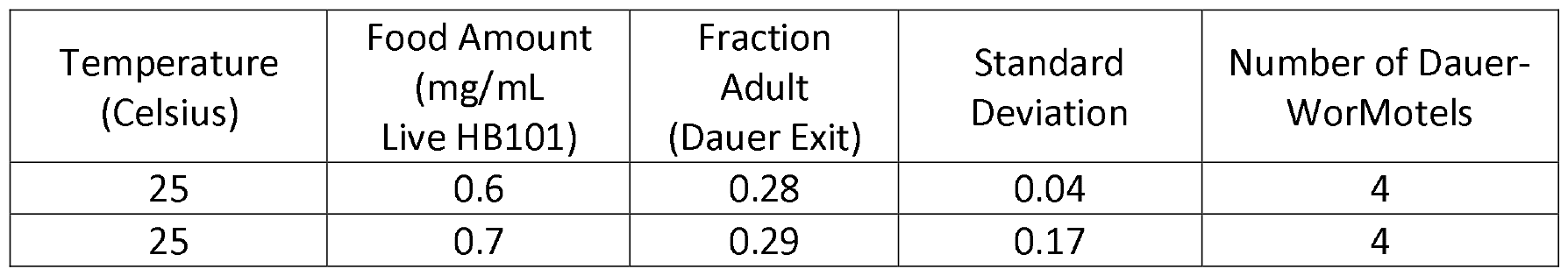
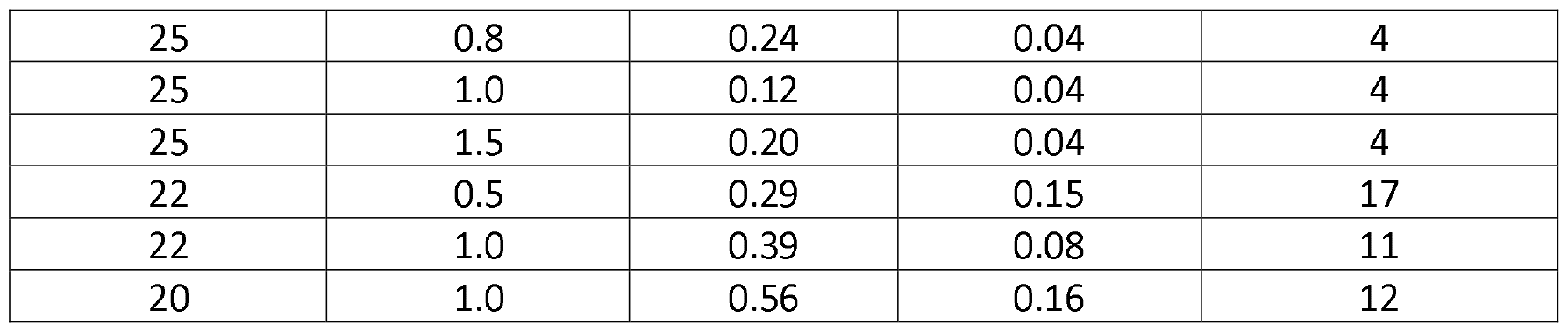
High temperature and low food (live HB101) decrease dauer exit. These data are from dauer exit assays in the Dauer-WorMotel with N2 animals. After starvation and SDS selection, dauer worms were individually housed in the listed food (live HB101 with 5 mg/mL cholesterol) and temperature for five days and then phenotyped. Each Dauer-WorMotel contains up to 48 dauer worms to start.

| Temperature (Celsius) | Food Amount (mg/mL Live HB101) | Fraction Adult (Dauer Exit) | Standard Deviation | Number of Dauer-WorMotels |
| --- | --- | --- | --- | --- |
| 25 | 0.6 | 0.28 | 0.04 | 4 |
| 25 | 0.7 | 0.29 | 0.17 | 4 |
| 25 | 0.8 | 0.24 | 0.04 | 4 |
| 25 | 1.0 | 0.12 | 0.04 | 4 |
| 25 | 1.5 | 0.20 | 0.04 | 4 |
| 22 | 0.5 | 0.29 | 0.15 | 17 |
| 22 | 1.0 | 0.39 | 0.08 | 11 |
| 20 | 1.0 | 0.56 | 0.16 | 12 |

### Dauer exit relies on IGF signaling from ASJ and neurons post-synaptic to ASJ

ASJ-triggered dauer exit is largely mediated by the release of insulin-like peptide INS-6, which binds DAF-2, which in turn promotes dauer exit (Hung *et al*., 2014). To confirm the role of INS-6 in dauer exit for our single-worm assay, we tested the *ins-4;ins-5;ins-6* triple mutant (ZM6523 [*hpDf761*]) in our Dauer-WorMotel and observed less dauer exit than N2, as expected (Figure 2A).

To block chemical neurotransmission from ASJ, tetanus toxin (TeTx) was expressed in the ASJ neuron under the *trx-1* promoter which is specific to ASJ. TeTx is a protease that cleaves synaptobrevin, a membrane protein important for synaptic vesicle release of neurotransmitters. We observed less dauer exit in the *Ptrx-1::TeTx::mCherry* line than in N2 (Figure 2B). Interestingly, no difference in dauer formation was observed previously between wildtype and *unc-13* mutants (synaptobrevin in *C. elegans*) (Shen *et al*., 2007). However, disrupting neurotransmission in the whole body may potentially mask antagonistic contributions of multiple neurons to the dauer-exit decision. When synaptic vesicle disruption was limited to only ASJ, less dauer-exit was observed, indicating synaptic transmission from ASJ promotes dauer exit.

While INS-6 is important for promoting dauer exit, multiple insulin-like peptides (ILPs) are expressed throughout the body of the worm in neuronal and non-neuronal tissues (Cornils, 2011; Zheng *et al*., 2018). DAF-2 is the sole IGFR in C. elegans, therefore it is the target of all ILPs. To test whether synaptic signaling from ASJ contributes to the dauer decision in addition to ILP signaling, we quantified dauer exit in a *daf-2 (e1370), Ptrx-1::TeTx::mCherry* background. We did not observe a difference in dauer exit between *daf-2* and *daf-2,trx-1::TeTx::mCherry* (Figure 2C). This indicates that while synaptic signaling form ASJ is important for dauer exit (Figure 2B), non-ILP signaling does not contribute significantly to the decision.

If dauer exit is primarily dependent on synaptic release of ILPs from ASJ, then blocking synaptic release in ASJ should produce similar probabilities of dauer exit as the *ins-4;ins-5;ins-6* triple mutant, ZM6523. While both ZM6523 and *Ptrx-1::TeTx::mCherry* decrease dauer exit (Figure 2A,B), blocking synaptic release in ZM6523 almost completely eliminated dauer exit (Figure 2D). In addition to ILPs, ASJ also releases the neurotransmitter acetylcholine, is post-synaptic to several neurons, and shares electrical synapses with the interneuron AIA (Cook *et al*., 2019). Since blocking synaptic release from ASJ in the *ins-4;ins-5;ins-6*, but not the *daf-2* (*e1370*) background, decreases dauer exit, it is likely that ILP release from neurons post-synaptic to ASJ also influence the dauer decision.

### Dauer exit relies on 12 hours of ASJ stimulation

ASJ normally responds to food odorants(citation). Odorants bind G-protein coupled receptors, which cause a signaling cascade that ends in the neuron’s depolarization (Cornelia I. Bargmann, 2006). However, the dynamic delivery of food to unrestrained dauers is challenging. As an alternative to food, we turned to optogenetics. One challenge with using channelrhodopsin over a long timescale is the prolonged exposure of animals to high intensities of light for long periods. However, a variant of channelrhodopsin (C128S) is sensitive to blue-light and does not require constant illumination to remain open (Berndt *et al*., 2009). Instead, blue light triggers the opening of the channel, which remains open for several minutes until closed by exposure to amber light. C128S has been used in *C. elegans* in the past to manipulate motor neurons and dauer exit over the hour timescale with low intensities of light (470 nm; <0.01 mW/mm^2^) (Schultheis *et al*., 2011). Our LED arrays produced light intensities in this range (Figure 3A). To quantify how the temporal dynamics of ASJ influence dauer exit, we depolarized ASJ with C128S at different intensities and time periods to measure how ASJ depolarization amplitude and duration influence dauer exit.

**Figure 3:**
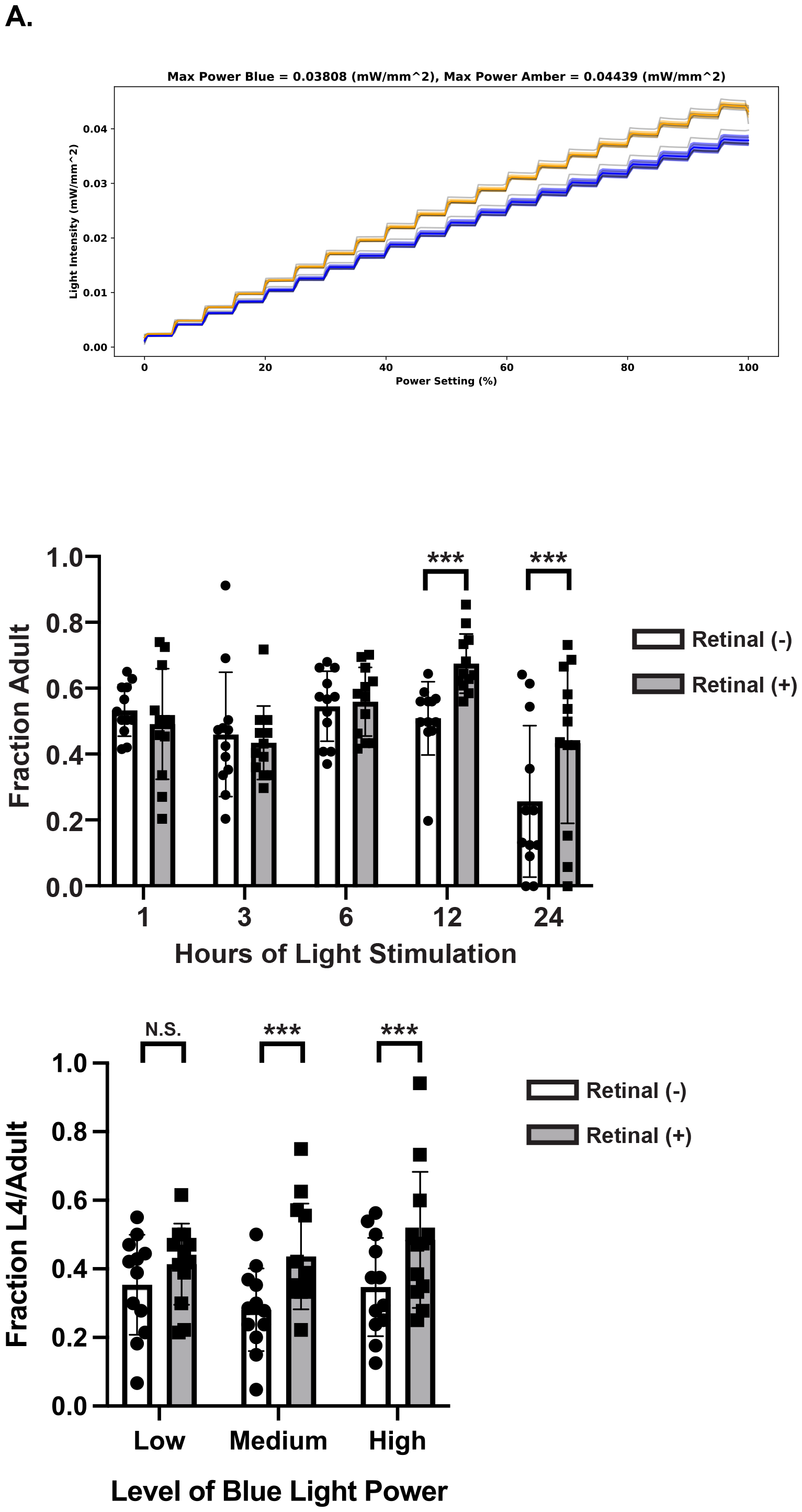
Optogenetic stimulation of ASJ drives dauer exit. **A**. Light intensities for blue (470 nm) and amber (590 nm) light in the optogenetics arena, measured at various power settings. **B**. The temporal window for ASJ activity to promoter dauer exit is 12 to 24 hours. *Ptrx-1::TeTx::mCherry* animals were stimulated at 470 nm: 8 µW/mm^2^ in the presence or absence of trans-retinal. The chi-square statistics are as follows: 24 hours - X^2^ (1, N = 337) = 174.0, p < 0.001 ; 12 hours - X^2^ (1, N = 409) = 198.0, p < 0.001 ; 6 hours - X^2^ (1, N = 437) = 213.0, p = 0.69; 3 hours - X^2^ (1, N = 379) = 181.0, p = 0.53 ; 1 hour - X^2^ (1, N = 389) = 201.0, p = 0.32. **C**. Dauer exit dependence on amplitude of ASJ photostimulation. *Ptrx-1::TeTx::mCherry* animals were stimulated in the presence or absence of trans-retinal for 12 hours under low (470 nm: 0.4 µW/mm^2^), medium (470 nm: 8 µW/mm^2^), and high (470 nm: 16 µW/mm^2^) light intensities. The chi-square statistics are the following: Low **-** X^2^ (1, N = 425) = 201.0, p = 0.15; Medium - X^2^ (1, N = 449) = 209.0, p < 0.001; High - X^2^ (1, N = 390) = 182.0, p < 0.001.

To define the temporal window of ASJ activity that is sufficient to trigger dauer exit, we stimulated *Ptrx-1::C128S:SL2::GFP* for different time intervals immediately after placing the dauers in the Dauer-WorMotel (Figure 3B). *Ptrx-1::C128S:SL2::GFP* worms were exposed to the following light conditions: a loop of 1 second 470 nm: 8 µW/mm^2^, followed by 5 seconds of darkness, followed by 2 seconds of 590 nm: 30 µW/mm^2^. This resulted in C128S depolarization for 6 seconds for every 8 second duty cycle. Since worms do not produce their own trans-retinal (the light absorbing co-factor for channelrhodopsin), worms were cultured with or without (control) trans-retinal in their food prior to developing into dauer animals. We found that photo-stimulation of ASJ for 6 hours or less was insufficient to increase dauer exit, but 12 hours or more was sufficient (Figure 3B).

In principle, increased depolarization of ASJ should suggest higher mean environmental quality. *Ptrx-1::C128S:SL2::GFP* animals were stimulated with the prior illumination protocol at low (470 nm: 0.4 µW/mm^2^), medium (470 nm: 8 µW/mm^2^), and high (470 nm: 16 µW/mm^2^) light intensities for 12 hours. We found statistically significant increased dauer exit at the medium and high illumination levels, as compared to the no-retinal controls (Figure 3C). We did not observe a difference at the low illumination level.

## Discussion

Here we present an experimental assay for dauer exit in *C. elegans* that is simple, scalable, and compatible with optogenetics. The illumination setup can easily fit within an incubator, and the manufacture of the devices simply requires casting PDMS. Animals can easily be phenotyped by observation under a dissecting microscope. The assay produces results under genetic perturbation that are consistent with the role of ILP signaling in dauer exit. Interestingly, blocking synaptic transmission in the absence of INS-6 further decreases dauer exit, indicating that other ILP-releasing neurons post-synaptic to ASJ contribute to the dauer exit decision. FLP-2 neuropeptides released by the interneuron AIA inhibit dauer entry, while INS-1 released by other interneurons also drive dauer entry (Chai *et al*, 2021). While these results did not address dauer exit, it indicates that the activity of interneurons that are post-synaptic to dauer-influencing sensory neurons also contribute to the dauer decision through neuropeptide signaling.

## Acknowledgements

We thank J. Kim, H. Zhao, E. Niebur, members of the Johns Hopkins University Biology and Neuroscience Departments and Gordus lab members for helpful discussions and comments on the manuscript. A.P. acknowledges funding from the Nathaniel Boggs, Jr. Ph.D. Scholar Fellowship. A.G. acknowledges funding from NIH (R35GM124883).

## Author contributions

A.P., J.M., A.C., A.M., I.M. and A.G. designed research. A.P. and J.M. performed all dauer experiments. A.P. designed the Dauer-WorMotel. A.C. designed the circuit board and chamber holders. A.P., A.C., and A.M. wrote software to control the LED boards. A.P., A.C., A.M. and I.M. assembled the circuit boards and holders. A.P. and A.G. wrote the manuscript.

